# scMAGS: Marker gene selection from scRNA-seq data for spatial transcriptomics studies

**DOI:** 10.1101/2022.03.22.485261

**Authors:** Yusuf Baran, Berat Doğan

## Abstract

Single-Cell RNA sequencing (scRNA-seq) has provided unprecedented opportunities for exploring gene expression and thus uncovering regulatory relationships between genes at the single cell level. However, scRNA-seq relies on isolating cells from tissues. Thus, the spatial context of the regulatory processes is lost. A recent technological innovation, spatial transcriptomics, allows to measure gene expression while preserving spatial information. A first step in the spatial transcriptomic analysis is to identify the cell type which requires a careful selection of cell-specific marker genes. For this purpose, currently scRNA-seq data is used to select limited number of marker genes from among all genes that distinguish cell types from each other. This study proposes scMAGS (**s**ingle-**c**ell **MA**rker **G**ene **S**election), a new approach for marker gene selection from scRNA-seq data for spatial transcriptomics studies. scMAGS uses a filtering step in which the candidate genes are extracted prior to the marker gene selection step. For the selection of marker genes, cluster validity indices, Silhouette index or Calinski-Harabasz index (for large datasets) are utilized. Experimental results showed that, in comparison to the existing methods, scMAGS is scalable, fast and accurate. Even for the large datasets with millions of cells, scMAGS could find the required number of marker genes in a reasonable amount of time with less memory requirements.

## Introduction

Organ systems consist of different cellular subpopulations. The location of these cellular subpopulations within the tissue is closely related to the function of the cells [1]. In recent years, scRNA-seq technology has allowed us to measure RNA levels (transcriptomics) of single cells which provides valuable insights into our understanding of cell types and functions. However, scRNA-seq experiments require isolation of single cells from tissue and this necessity leads to the loss of the spatial position of the cells in the tissue and their spatial proximity to each other. Recent studies show that cell-cell communication (CCC) regulated by biochemical signals is an important aspect of tissue structure and function, regulates individual cellular processes and is effective in intercellular interaction [2]. Knowing the spatial positions of cells in the tissue is critical to understanding normal development and disease pathology. Spatial transcriptomics offers a solution to this problem by providing information about the location of cell types identified by the mRNA readouts in tissue. By combining spatial information with scRNA-seq data, several methods have been developed to measure gene activity in a tissue sample and to map where the activity is occurring [3-10].

Current spatial transcriptomics technologies are primarily categorized as sequencing-based and imaging-based methods [11-17]. Fundamental aspects of these spatial transcriptomics technologies can vary widely, both in terms of the number of genes that can be probed and the size of tissue that can be assayed [18]. For instance, the number of genes that can be probed in single-molecule fluorescence in situ hybridization (smFISH), one of the imaging-based methods, is experimentally constrained by the product of the number of fluorescent channels and the hybridization cycles [19]. Currently, state-of-the-art smFISH methods use around 40 genes [20]. Thus, using labeled scRNA-seq data to optimally select a limited number of marker genes that distinguish cell types from among all genes is a combinatorially difficult problem.

In literature, recently a number methods have been proposed for selection of marker genes from scRNA-seq data to distinguish cell types from each other. One of these methods is the COMET [21] in which k marker genes are selected and sorted by one-vs-all methodology. However, the complexity of this method is G^k^, where G represents the number of genes. Thus, for even k = 4, it becomes quite infeasible to select markers. Indeed, the COMET provides at most 4 marker genes. In addition, one-vs-all approach cannot offer a solution to the problem when the number of cell types is greater than the number of markers [22]. In a recent study, authors proposed the scGeneFit for optimal selection of marker genes [20]. scGeneFit selects gene markers that jointly optimize cell label recovery using label-aware compressive classification methods. However, this method only focuses on the selection of markers that correctly separates cells with different labels from each other. This may lead a high classification accuracy with the selected subset of markers but not necessarily provide the ideal set of marker genes for spatial transcriptomics studies. An ideal marker gene must be highly expressed in a certain cell type and lowly expressed (ideally zero expression) in all other cell types. A recent study [23] showed that, scGeneFit selects markers some of which are highly expressed in a number of cell types which is not suitable for spatial transcriptomics studies. Selecting the proper marker genes require a careful filtering process. In [23], authors filter out the lowly and highly expressed genes so that only the genes those are expressed in greater than 30% of the classes of interest and in less than 75% of cells with more than 50% of the classes of interest are retained. The filtered genes are then ranked according to an ensemble learning model or a deep neural network, generating a final set of markers for each cell type. The overall framework is called as SMaSH (Scalable Marker (gene) Signal Hunter) which is publicly available as a python package. Although, SMaSH could select appropriate marker genes for some datasets, it has still some problems for most of the datasets. SMaSH rely on supervised selection of marker genes by a number of classification methods. Therefore, in cases where the number of samples is not sufficient for a certain type of cell, the model used in the training phase of SMaSH could fail to provide the appropriate markers. Additionally, as in the scGeneFit, a high classification accuracy does not necessarily mean the algorithm provides the appropriate markers for the spatial transcriptomics studies. In cases where a specific gene is differentially expressed among different cell types is not a challenge for nonlinear classification algorithms to provide a high classification accuracy. Indeed, the gene filtering approach used in the SMaSH lacks foresight for such scenarios. Therefore, for a number of datasets, SMaSH selects marker genes that are expressed in a number of cell types which is not appropriate for spatial transcriptomics studies. Another shortcoming of the SMaSH package is that it cannot work with sparse matrices. Recently, scRNA-seq experiments are conducted for millions of cells and the generated massive data need to be stored in more efficient formats such as the sparse matrix format. This shortcoming limits the usability of the method for large-scale scRNA-seq data. In a more recent study, authors proposed the COSine similarity-based marker Gene identification (COSG), to select the cell specific marker genes from scRNA-seq, scATAC-seq and spatially resolved transcriptomics data [24]. In comparison to the previous studies, COSG is fast and scalable for ultra-large datasets of million-scale cells. However, it has a significant memory requirement and for some cases the cell-specific marker genes found by the COSG are shown to be expressed not only in the target cell type but also in other cell types.

This study proposes scMAGS (**s**ingle-**c**ell **MA**rker **G**ene **S**election), a new approach for marker gene selection for spatial transcriptomics studies. scMAGS uses a filtering step in which the candidate genes are extracted prior to the marker gene selection step. For the selection of marker genes, cluster validity indices, Silhouette index or Calinski-Harabasz index (for large datasets) are utilized. Comparison with the existing methods indicated that, scMAGS selects markers genes that are exclusive to each cell type such that, the corresponding markers genes are highly expressed in a specific cell type while lowly expressed (either zero expression) in other cell types which is desired for the spatial transcriptomics studies. Moreover, experiments showed that the scMAGS is computationally efficient and requires less memory and execution time. Even for the large datasets with millions of cells, scMAGS could find the marker genes in a reasonable amount of time with less memory requirements.

## Materials and Methods

### Datasets

The input of the scMAGS is a scRNA-seq expression data with cells in rows and genes in columns. Expression data in a sparse matrix form is also allowed. The cell labels must be provided as a separate input.

Table S1 lists the details of nineteen datasets used to evaluate the performance of the method. All datasets were obtained from publicly available databases.

### Filtering candidate genes from the log-transformed scRNA-seq data

Gene filtering is one of the most important features that distinguishes the scMAGS from other methods. When selecting marker genes for each cell type, it is not necessary to search for all genes. For the large scale scRNA-seq datasets, the execution of marker selection with all genes can cause computational difficulties and requires high memory. scMAGS identifies candidate marker genes for all cell types with a cluster-specific gene filtering step, and marker selection is carried out on the identified candidate genes. This reduces the memory usage and thus the computational cost.

The following criteria should be considered when filtering the marker genes for spatial transcriptomic probes: i) High expression rate in the specific cell type ii) Low (ideally zero) expression rate in the other cell types iii) High average expression value in the specific cell type iv) Low average expression value (ideally zero) in other cell types.

A classic scRNA-seq experiment usually contains more than 20,000 genes. Elimination of genes that do not differentiate between cell types is crucial for the efficiency and scalability of the algorithm. A typical drop-based (drop-seq) scRNA-seq data may contain up to 90% zero values in the expression matrix [25]. The high number of genes can be greatly reduced by filtering out genes that are not expressed in more than a few cells or are expressed at the same levels in all cells, thus do not provide information about cellular heterogeneity [26].

scMAGS first calculates the expression rates of all genes in all cell types. As a preprocessing step prior to filtering, it removes genes with less than 20% expression in all cell types. After eliminating genes that do not inform about cellular heterogeneity, the next step is normalization. Count values are converted to expression levels by the normalization process. The count matrix should be normalized to reduce the bias of experimental effects [27]. The number of reads for a gene in each cell is expected to be proportional to the gene-specific expression level and cell-specific scaling factors [28]. However, even if the cells are identical, the count depths may be different for each cell. Difficulty in preparing libraries from cells containing minimal mRNA material, technical bias in cDNA capture or PCR amplification can cause this problem.

The normalization step scales the count data to obtain accurate gene expression values across cells, in other words, to bring all cells under the same conditions [28]. Without normalization, incorrect marker selection can be made due to technical biases. Therefore, before gene filtering, log(1+x) transform is applied to the entire count matrix. log(1+x) transform reduces the skewness in data.

Let *X* ∈ *R*^*nxm*^, *X* = {*X*_1_, *X*_2_, …, *X*_*k*_} be the resulting preprocessed and log(1+x) transformed expression matrix with n cells and m genes. *X* could be considered as a combination of *k* matrices where *k* represents the number of different cell types or states. Each cell type or state could be considered as a separate cluster. For computational efficiency, gene filtering is performed for each cluster separately and then the marker gene selection process is carried out on these candidate genes for each cluster. For cluster-specific gene filtering, firstly, within-cluster expression rates and within-cluster average expression values of all genes are computed over the *X* matrix. Let *A* ∈ *R*^*kxm*^ be the expression rate matrix computed from *X*. For each gene at each cluster, the expression rate is computed by dividing the number of cells with non-zero expression to the total number of cells within that particular cluster. Let *B* ∈ *R*^*kxm*^ be the average expression matrix computed from *X*. For each gene at each cluster, the average expression value is computed by averaging the expression values of that particular gene at that particular cluster. With the help of *A* and *B* matrices, cluster-specific candidate marker genes are found by following the below steps for each cluster *k*. A step-by-step description of the candidate marker gene filtering process is also shown in Figure 1.

**Figure 1.**
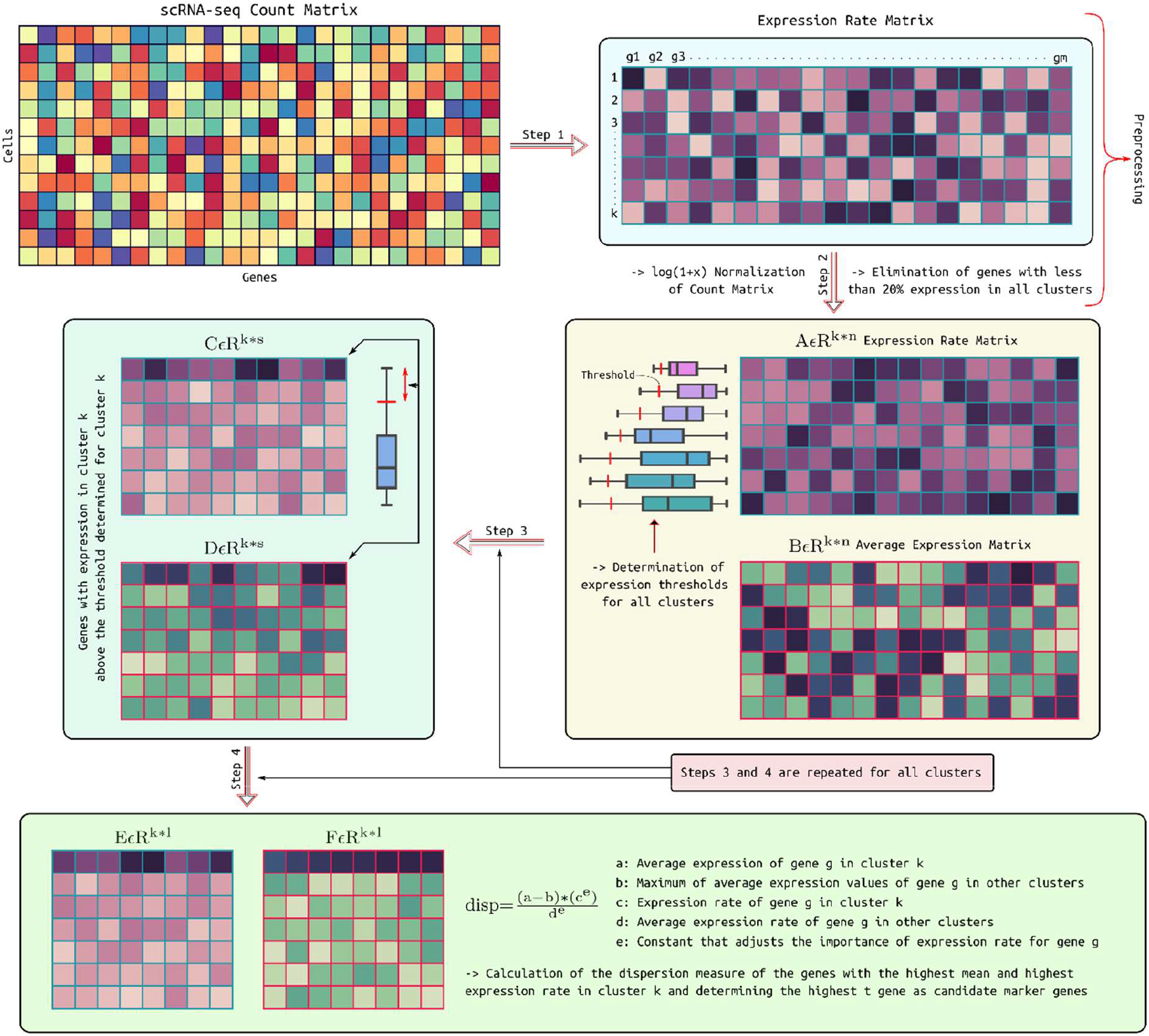
A step-by-step description of the gene filtering process providing the candidate marker genes for gene selection step. Each four steps are repeated for each cluster to find the cluster-specific candidate marker genes.

1. By using the matrix *A*, for cluster *k*, find the genes those have an expression rate above a threshold. For each cluster *k*, the threshold is calculated with Eq.1. In Eq.1, *Q*_3_ represents the 75^th^ percentile and *IQR* represents the interquartile range (difference between 75^th^ and 25^th^ percentiles).

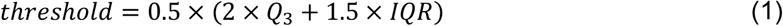
2. For cluster k, form the matrix *C* ∈ *R*^*kxs*^ from matrix *A* and form the matrix *D* ∈ *R*^*kxs*^ from matrix *B*, by using the genes those have an expression rate above threshold. Here, s represents the number of genes that are above the threshold.
3. From matrix *D*, find the genes with maximum average expression in cluster *k*, and form the matrix *F* ∈ *R*^*kxl*^. Here, *l* represents the number of genes with maximum average expression in cluster k. By using the same genes, form the matrix *E* ∈ *R*^*kxl*^ from matrix *C*.
4. For each gene *g*, compute a dispersion metric. To do this, by using matrices *E* and *F* find the following constants.

a. average expression of gene *g* in cluster *k* (from matrix *F*)
b. maximum of average expression values of gene *g* in other clusters (from matrix *F*)
c. expression rate of gene *g* in cluster *k* (from matrix *E*)
d. average expression rate of gene *g* in other clusters (from matrix *E*)
e. constant that adjusts the importance of expression rate for gene *g*

By using the above defined constant values, for each gene *g* the dispersion metric is computed as in Eq.2. Then, genes are sorted according to their dispersion values and top *t* genes with the highest dispersion values are selected as the candidate marker genes of cluster *k*.

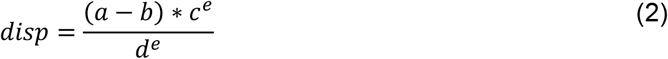

### Selecting marker genes from the set of filtered candidate genes

To select the marker genes from the cluster-specific candidate marker gene set, either Silhouette index or for large datasets (>100.000 cells), Calinski-Harabasz index is used. The Silhouette index is a measure of how similar a sample is to its own cluster compared to other clusters and it has been previously shown to be the most efficient method to evaluate the clustering validity among thirty different validity indices [29]. The Silhouette index ranges from −1 to +1, where a high value indicates that the object is well matched to its own cluster and poorly matched to neighboring clusters. On the other hand, the Calinski-Harabasz index, is the ratio of between-cluster and within-cluster distribution. The score is higher when the clusters are dense and well separated from each other. Because its computationally efficient, it is preferred over Silhouette index in large datasets. Moreover, it was shown to be the second most efficient method to evaluate the clustering validity [29]. Considering the marker gene selection problem, the set of genes that leads to the optimal partition of cells is expected to maximize both the Silhouette or Calinski-Harabasz scores of overall data.

#### Silhouette index

To obtain the silhouette score for an object *o*_*i*_, first the within-cluster average distance *a*(*i*) between *o*_*i*_ ∈ *C*_*k*_ and all other objects *o*_*i*′_ in cluster *C*_*k*_ is computed as in Eq.3. In Eq.3 *n*_*k*_ represents the number of objects in cluster *k* and *d*(*o*_*i*_, *o*_*i*′_) represents the Euclidean distance between the object *o*_*i*_ and *o*_*i*′_.

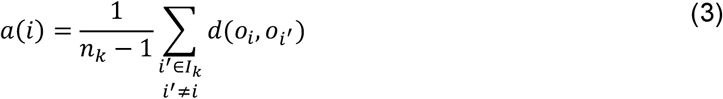

Then, the average distances *δ*(*o*_*i*_, *C*_*k*′_) between *o*_*i*_ and all other objects in other clusters *C*_*k*′_ is computed as in Eq.4.

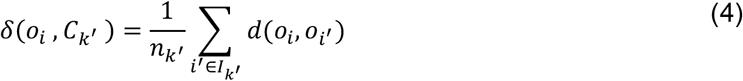

Let *b*(*i*) be the smallest of these average distances:

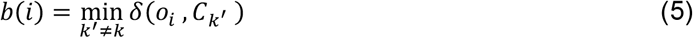

For each object *o*_*i*_, the Silhouette score *s*(*i*) is then computed as in Eq.6.

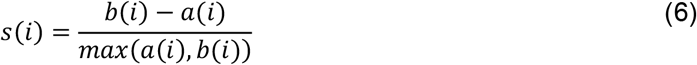

The average of Silhouette score for a given cluster *C*_*k*_ is called as the cluster mean Silhouette and is computed as in Eq.7.

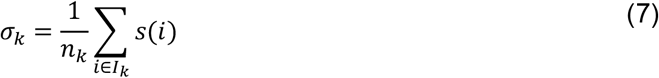

Finally, the global Silhouette index of the data is computed by averaging the mean Silhouette score of all clusters as in Eq.8.

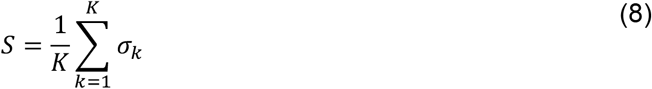

#### Calinski-Harabasz index

To obtain the Calinski-Harabasz score of a labeled dataset containing *k* clusters, first the between-cluster scattering matrix *B*_*k*_ and within-cluster scattering matrix *W*_*k*_ should be computed. The matrices *B*_*k*_ and *W*_*k*_ can be computed with Eqs. 9-10.

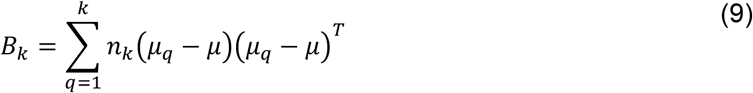

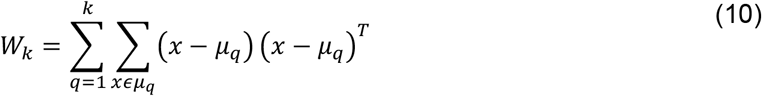

In Eqs.9-10, *k* represents the number of clusters, *n*_*k*_ represents the number of samples in cluster *q, µ*_*q*_ represents the center of the cluster *q* and *µ* represents the center of overall data. Having computed the matrices *B* and *W*, the Calinski-Harabasz score of a dataset can be computed as in Eq.11.

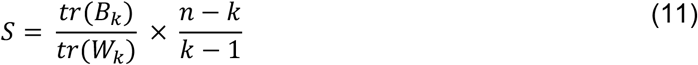

In Eq.11, *n* represents the number of samples in the dataset.

## Results

The performance of the scMAGS is compared to the existing methods scGeneFit, SMaSH and COSG by using the datasets listed in Table S1. The methods scGeneFit and SMaSH were only run for some of the datasets either because the method required more computational resources or it failed to run because of an algorithmic problem. All experiments were carried out on a Dell Precision T7820 workstation equipped with two Intel Xeon® Silver 4210R 2.40GHz 20 core processors and 128 GB RAM.

### Peak memory requirements and computation time

To evaluate memory usage and computation time, all methods were run with their default parameters to select three marker genes for each cell type. The scGeneFit and SMaSH are not able to run with sparce matrices. Therefore, for some of the datasets, the sparce matrices were converted into dense matrices for these two methods. However, for the large datasets the conversion process requires extensive memory requirements (> 400GB RAM) and therefore, it was not possible to evaluate the performance of these methods for the large datasets. In addition, the SMaSH uses classification algorithms which need partitioning of a dataset into training and test sets. However, for some of the datasets, this leads the method to fail during the execution when the size of the test set for a certain cell type is not sufficient to keep the algorithm running. Therefore, the method was not able to run for most of the datasets listed in Table S1.

On the other hand, because of the high memory requirements, scGeneFit was not able to run with datasets with high number of genes, independent of number of cells. Even if it worked with a dataset, it exhibited the worst performance in terms of computational time. In Table S2, the computation time required to run each method is listed for each dataset. As shown in Table S2, scMAGS and COSG were able to run with all datasets in a quite reasonable time. Although, the COSG is much faster for most of the datasets, the results provided by scMAGS is comparable and it takes only 4.8 minutes to run Bhaduri dataset with 1.3M cells, for which COSG needs 4.6 minutes to find the marker genes. On the other hand, considering the peak memory requirements of the methods (Table S3), the scMAGS performs best and overcomes the COSG with a peak memory requirement of 46949 MB for the Bhaduri dataset which is much smaller than the one required by COSG, 112249 MB.

### Expression profiles of selected marker genes

Selected marker genes by each method are evaluated based on the criteria provided earlier. A marker gene must have a high expression rate within a specific cell type with an average high expression value. Although, the differential expression of a gene in different cell types provides a high classification accuracy, this does not necessarily mean it is a good candidate to be considered as a marker gene. This fact is shown from the confusion matrices (Figure 2) and dotplots [30] (Figure 3) obtained for each method with the Zeisel dataset, which is one of the datasets that the four methods were able to run commonly. As shown in Figure 2, the k-Nearest Neighbor (k-NN) classification results obtained by using the selected marker genes for each method do not exhibit a significant difference. However, the dotplots shown in Figure 3 suggests that the genes selected by the scGeneFit and SMaSH methods are highly expressed in more than one cell type which make them poor candidates to be considered as marker genes. On the other hand, the genes selected by the scMAGS and COSG have high expression rate and high average expression values in a specific cell type with low expression rates and low average expression values in other cell types which makes them ideal candidates to be considered as marker genes. This fact was further verified with another dataset that was commonly used to evaluate the performance of the four methods. In Figure 4, the dotplots obtained for the Kleshchevnikov dataset are shown. From this figure, it is obvious that, the scGeneFit and SMaSH methods again fails to find the proper marker genes. Although, some of the genes selected by scMAGS and COSG have also expression in other cell types, in general the selected genes meet the criteria to be considered as marker genes.

**Figure 2.**
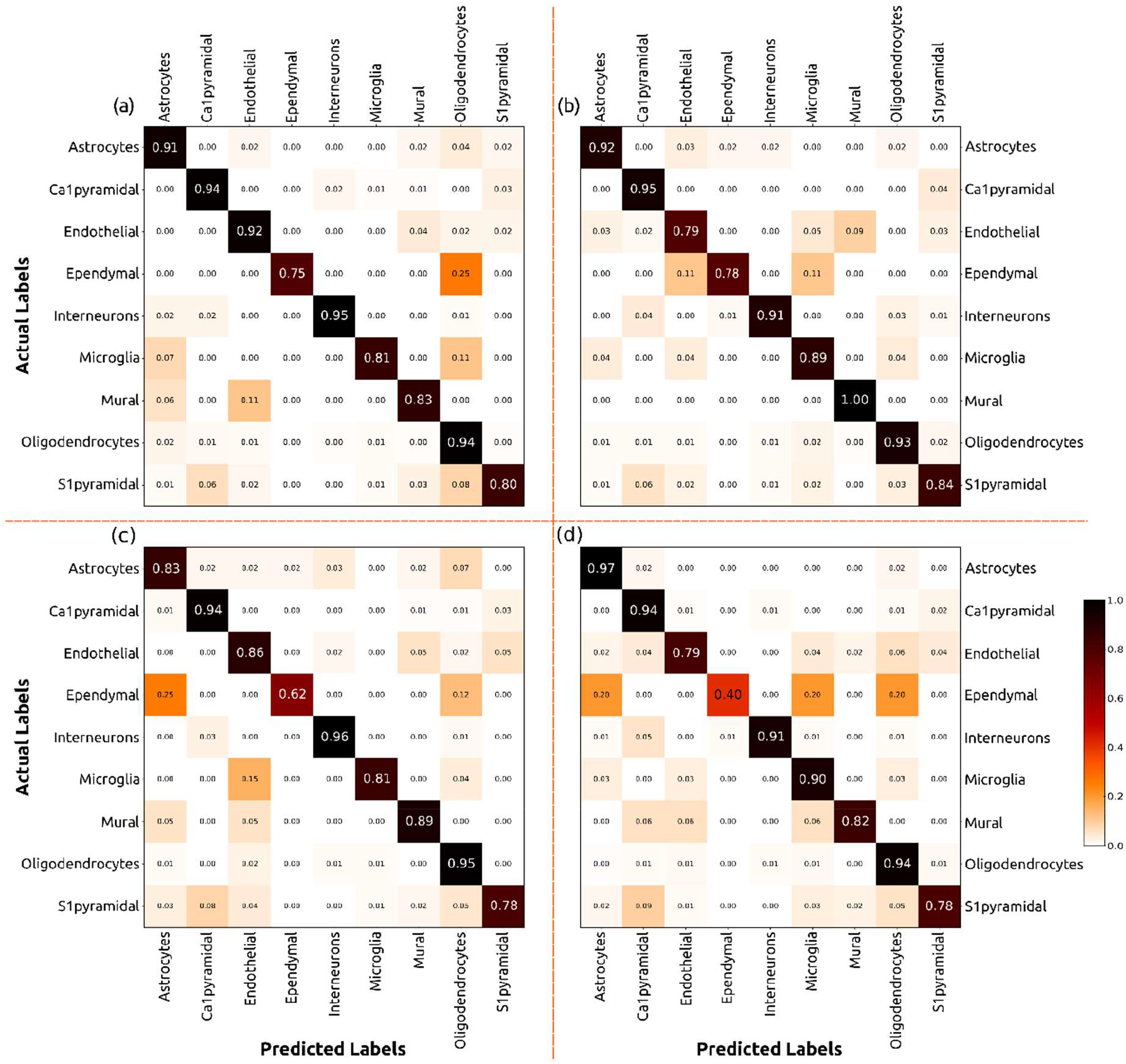
Confusion matrices of the k-Nearest Neighbor (k-NN) classification results obtained by using marker genes found for the Zeisel dataset **(a)** scMAGS **(b)** SMaSH **(c)** scGeneFit **(d)** COSG

**Figure 3.**
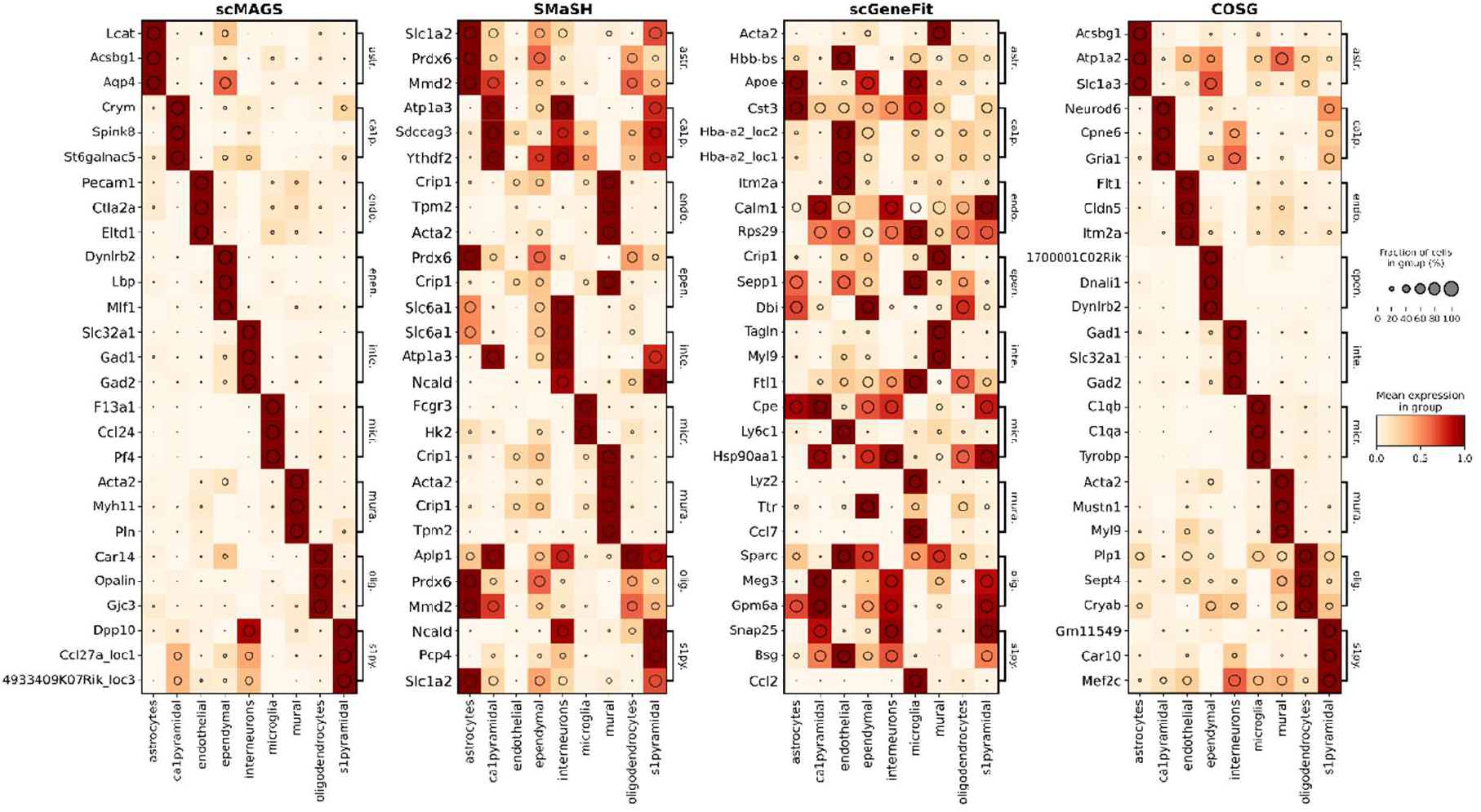
Dotplots of the marker genes found for the Zeisel dataset. Marker genes found by the scGeneFit and SMaSH methods are highly expressed in more than one cell type which make them poor candidates to be considered as marker genes.

**Figure 4.**
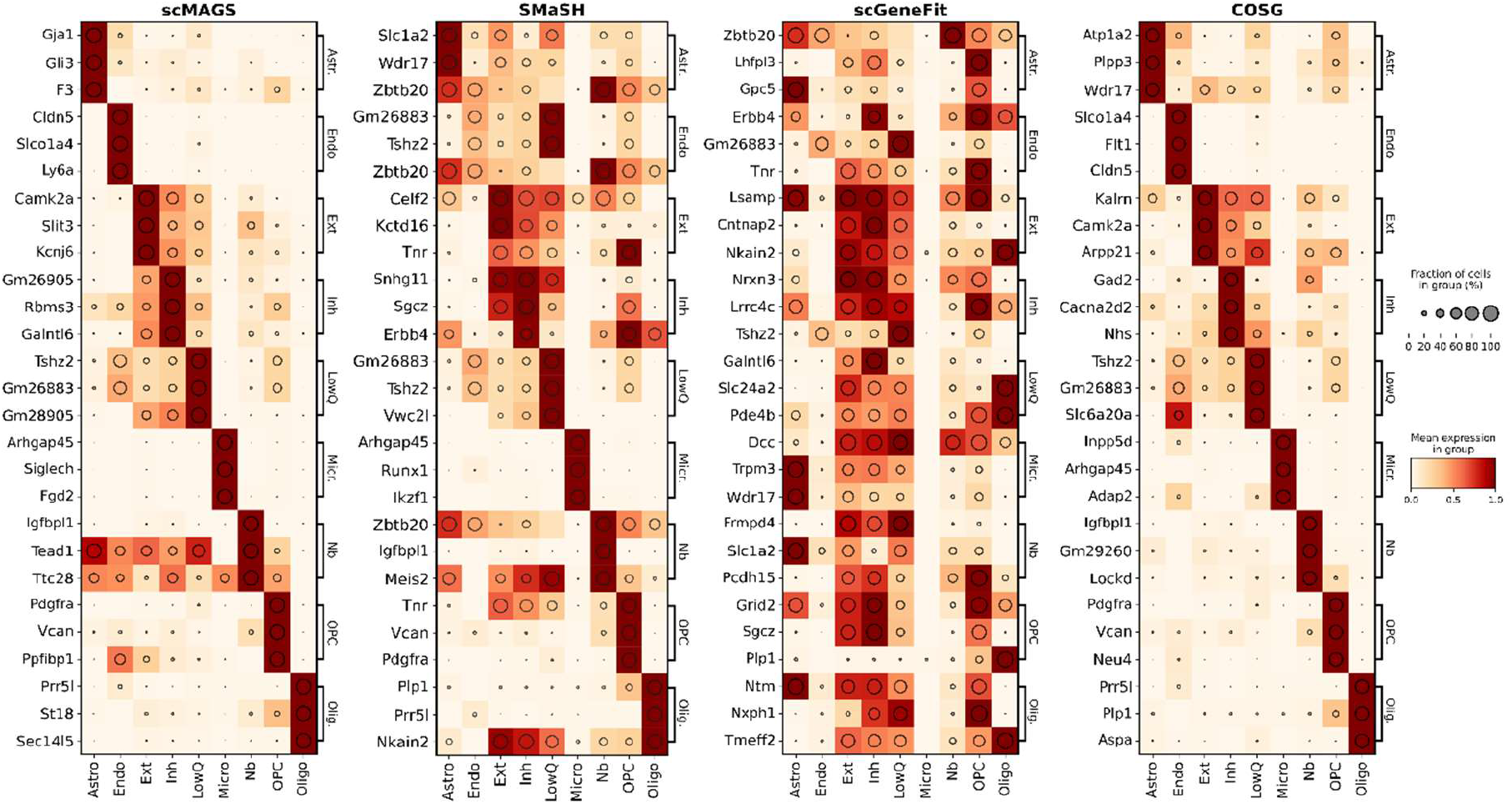
Dotplots of the marker genes found for the Kleshchevnikov dataset. Marker genes found by the scGeneFit and SMaSH methods are highly expressed in more than one cell type which make them poor candidates to be considered as marker genes.

A comparative analysis of the scMAGS and COSG for the other datasets are also provided in Figures S1-S5. As shown from these dotplots, for most of the datasets cell specific marker genes selected by COSG are expressed in a number of cell types which is not desired. The dotplot is one of the standard plots used in different studies to compare the expression rates and average expression values [23-24]. However, because it scales the expression rate and average expression of a gene to [0,1] interval among the different cell types, it might be misleading to solely rely on this plot. Therefore, for each dataset, distribution of the expression rates for the marker genes selected by the scMAGS and COSG methods are also compared to see whether there is a statistically significant difference between two groups.

In Figure 5, it is clear that, the marker genes selected by scMAGS have high within-cell expression rates and have low expression rates in other cell types. Note that, as in the scMAGS, for some of the datasets the marker genes selected by COSG also have high within-cell expression rates. On the other hand, for a number of datasets the interquartile range of boxplots is quite large, which means COSG selects marker genes not only with high within-cell expression rates, but genes with low within-cell expression rates are also selected. Nonetheless, for within-cell expression rates of the marker genes, in general, there is not any statistically significant difference (p > 0.05) between the results obtained by the scMAGS. However, the expression rates of the marker genes selected by COSG are also high in other cell types and, in comparison to scMAGS, for this case the difference is statistically significant (p < 0.05) for most of the datasets. This problem is more obvious and could be better investigated from the t-SNE plots and heatmaps shown in Figure 6 for one of the datasets (Baron Human 2 dataset). In this figure, even the best marker genes (first column of the heatmap for each cell type) selected by COSG for alpha and beta cells have expression in almost all other cell types.

**Figure 5.**
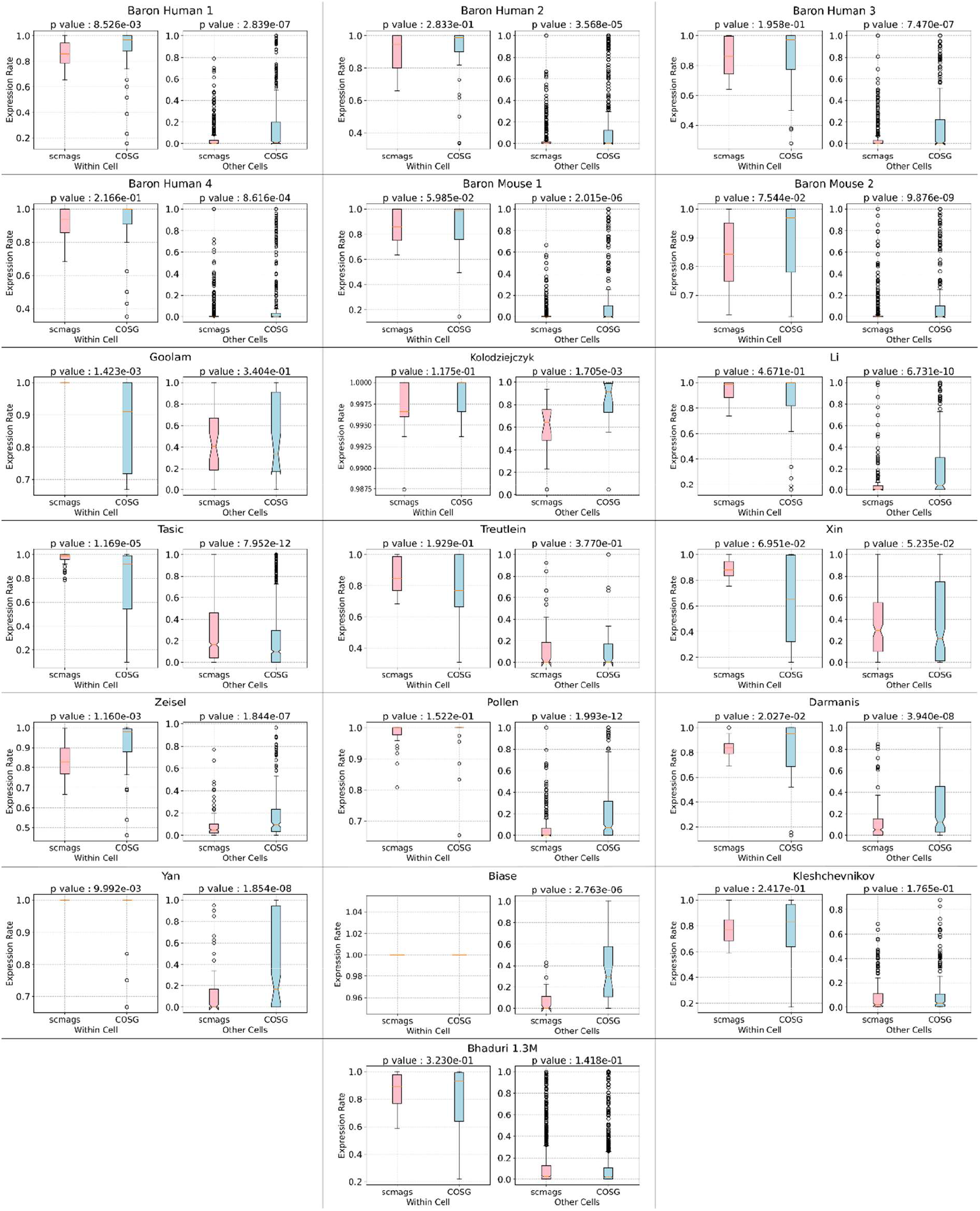
Marker genes selected by the scMAGS and COSG methods are evaluated by statistical comparison of their within-cell expression rates and expression rates in other cell types. A Mann-Whitney U test along with the box-whisker plots of the expression rates validated that, the marker genes selected by the COSG method have high expression rates in other cell types which is not desired.

**Figure 6.**
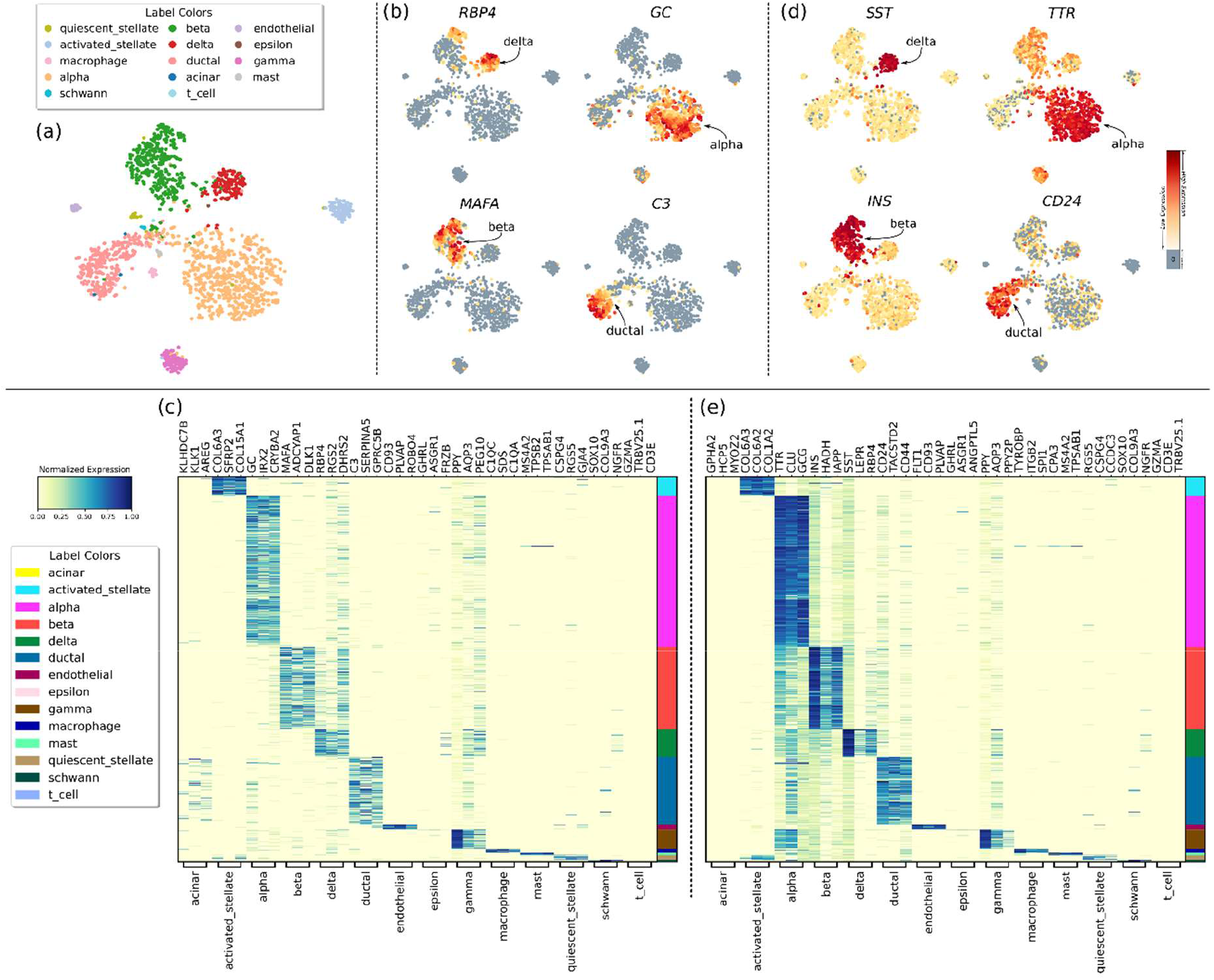
**(a)** Two-dimensional t-SNE plot of the Baron Human 2 dataset. **(b)** The expression of the best marker gene selected by the scMAGS method at each cell type **(c)** A heatmap of the marker genes selected by the scMAGS method **(d)** The expression of the best marker gene selected by the COSG method at each cell type **(e)** A heatmap of the marker genes selected by the COSG method

## Discussion

Understanding the spatial distribution of gene expression has become possible with the recently developed technologies. This has contributed to the solution of many fundamental problems of developmental biology. A major step in determining the spatial distribution of the gene expression across various tissues is the selection of accurate marker genes. Here, we present the scMAGS as a scalable, fast and accurate method for the marker gene selection from scRNA-seq data. scMAGS is implemented in Python and is made publicly available at GitHub.

The main difference of the scMAGS from the previous studies is its accurate gene filtering step by which the optimal set of candidate genes are extracted from the large-scale scRNA-seq data. The filtering step positively affects the memory requirement of the method and makes it computationally lightweight in comparison to the available methods. Next, by utilizing cluster validity indices (Silhouette and Calinski-Harabasz), scMAGS selects the ideal marker genes from the set of filtered candidate genes. The recently proposed COSG method is fast in comparison to the scMAGS. However, it requires more memory and for most of the datasets it selects genes that are not only expressed in the target cell type but also in other cell types which is not desired for spatial transcriptomics studies. Moreover, for a number of datasets, some of the marker genes selected by the COSG have low within-cell expression rates.

By default, the scMAGS selects three marker genes. However, this number can be changed based on the user requirements. Selected marker genes can be visualized by a number of plots such as the dotplot, heatmap, t-SNE distribution of marker genes in 2D and the confusion matrix obtained by the k-NN classification result.

In conclusion, the unique features of the scMAGS method in comparison to the previously published studies makes it a good alternative for marker gene selection problem. With its outstanding performance, it can serve as a general tool not only for spatial transcriptomics studies but also in other biological studies that require the accurate and fast identification of marker genes.

## Code availability

The scMAGS is made freely available as a Python package. A CLI (Command Line Interface) has been built internally in the package, so it can be run from the terminal environment without the need for any IDE (Integrated Development Environment). The source code of the scMAGS package is freely available at https://github.com/doganlab/scmags and can be downloaded from the Python Package Directory (PyPI) software repository with the command *pip install scmags*.

## References

1. Longo SK, Guo MG, Ji AL, Khavari PA. Integrating single-cell and spatial transcriptomics to elucidate intercellular tissue dynamics. Nat Rev Genet. 2021;22(10):627–644. Doi:10.1038/s41576-021-00370-8

2. Almet AA, Cang Z, Jin S, Nie Q. The landscape of cell-cell communication through single-cell transcriptomics. 15urrO pin Syst Biol. 2021;26:12–23. Doi:10.1016/j.coisb.2021.03.007

3. Zhu Q, Shah S, Dries R, Cai L, Yuan GC. Identification of spatially associated subpopulations by combining scRNAseq and sequential fluorescence in situ hybridization data. Nat Biotechnol. 2018;36(12):1183–1190. Doi:10.1038/nbt.4260

4. Karaiskos N, Wahle P, Alles J, et al. The Drosophila embryo at single-cell transcriptome resolution. Science. 2017;358(6360):194–199. Doi:10.1126/science.aan3235

5. Bageritz J, Willnow P, Valentini E, Leible S, Boutros M, Teleman AA. Gene expression atlas of a developing tissue by single cell expression correlation analysis. Nat Methods. 2019;16(8):750–756. Doi:10.1038/s41592-019-0492-x

6. Achim K, Pettit JB, Saraiva LR, et al. High-throughput spatial mapping of single-cell RNA-seq data to tissue of origin. Nat Biotechnol. 2015;33(5):503–509. Doi:10.1038/nbt.3209

7. Satija R, Farrell JA, Gennert D, Schier AF, Regev A. Spatial reconstruction of single-cell gene expression data. Nat Biotechnol. 2015;33(5):495–502. Doi:10.1038/nbt.3192

8. Halpern KB, Shenhav R, Matcovitch-Natan O, et al. Single-cell spatial reconstruction reveals global division of labour in the mammalian liver. Nature. 2017;542(7641):352–356. Doi:10.1038/nature21065

9. Butler A, Hoffman P, Smibert P, Papalexi E, Satija R. Integrating single-cell transcriptomic data across different conditions, technologies, and species. Nat Biotechnol. 2018;36(5):411–420. Doi:10.1038/nbt.4096

10. Stuart T, Butler A, Hoffman P, et al. Comprehensive integration of single-cell data. Cell. 2019;177(7):1888-1902.e21. doi:10.1016/j.cell.2019.05.031

11. Zhuang X. Spatially resolved single-cell genomics and transcriptomics by imaging. Nat Methods. 2021;18(1):18–22. Doi:10.1038/s41592-020-01037-8

12. Larsson L, Frisén J, Lundeberg J. Spatially resolved transcriptomics adds a new dimension to genomics. Nat Methods. 2021;18(1):15–18. Doi:10.1038/s41592-020-01038-7

13. Crosetto N, Bienko M, van Oudenaarden A. Spatially resolved transcriptomics and beyond. Nat Rev Genet. 2015;16(1):57–66. Doi:10.1038/nrg3832

14. Moor AE, Itzkovitz S. Spatial transcriptomics: paving the way for tissue-level systems biology. 16urrO pin Biotechnol. 2017;46:126–133. Doi:10.1016/j.copbio.2017.02.004

15. Asp M, Bergenstråhle J, Lundeberg J. Spatially resolved transcriptomes-next generation tools for tissue exploration. Bioessays. 2020;42(10):e1900221. Doi:10.1002/bies.201900221

16. Waylen LN, Nim HT, Martelotto LG, Ramialison M. From whole-mount to single-cell spatial assessment of gene expression in 3D. Commun Biol. 2020;3(1):602. Doi:10.1038/s42003-020-01341-1

17. Teves JM, Won KJ. Mapping cellular coordinates through advances in spatial transcriptomics technology. Mol Cells. 2020;43(7):591–599. Doi:10.14348/molcells.2020.0020

18. Rao A, Barkley D, França GS, Yanai I. Exploring tissue architecture using spatial transcriptomics. Nature. 2021;596(7871):211–220. Doi:10.1038/s41586-021-03634-9

19. Codeluppi S, Borm LE, Zeisel A, et al. Spatial organization of the somatosensory cortex revealed by cyclic smFISH. bioRxiv. Published online 2018. Doi:10.1101/276097

20. Dumitrascu B, Villar S, Mixon DG, Engelhardt BE. Optimal marker gene selection for cell type discrimination in single cell analyses. Nat Commun. 2021;12(1):1186. Doi:10.1038/s41467-021-21453-4

21. Delaney C, Schnell A, Cammarata LV, et al. Combinatorial prediction of marker panels from single-cell transcriptomic data. Mol Syst Biol. 2019;15(10):e9005. Doi:10.15252/msb.20199005

22. Reboredo H, Renna F, Calderbank R, Rodrigues MRD. Bounds on the number of measurements for reliable compressive classification. IEEE Trans Signal Process. 2016;64(22):5778–5793. Doi:10.1109/tsp.2016.2599496

23. Nelson ME, Riva SG, Cvejic A. SMaSH: A scalable, general marker gene identification framework for single-cell RNA sequencing and Spatial Transcriptomics. bioRxiv. Published online 2021. Doi:10.1101/2021.04.08.438978

24. Dai M, Pei X, Wang XJ. Accurate and fast cell marker gene identification with COSG. Brief Bioinform. Published online 2022. Doi:10.1093/bib/bbab579

25. Klein AM, Mazutis L, Akartuna I, et al. Droplet barcoding for single-cell transcriptomics applied to embryonic stem cells. Cell. 2015;161(5):1187–1201. Doi:10.1016/j.cell.2015.04.044

26. Zheng GXY, Terry JM, Belgrader P, et al. Massively parallel digital transcriptional profiling of single cells. Nat Commun. 2017;8(1):14049. Doi:10.1038/ncomms14049

27. Luecken MD, Theis FJ. Current best practices in single-cell RNA-seq analysis: a tutorial. Mol Syst Biol. 2019;15(6):e8746. Doi:10.15252/msb.20188746

28. Cao J, Spielmann M, Qiu X, et al. The single-cell transcriptional landscape of mammalian organogenesis. Nature. 2019;566(7745):496–502. Doi:10.1038/s41586-019-0969-x

29. Arbelaitz O, Gurrutxaga I, Muguerza J, Pérez JM, Perona I. An extensive comparative study of cluster validity indices. Pattern Recognit. 2013;46(1):243–256. Doi:10.1016/j.patcog.2012.07.021

30. Wolf FA, Angerer P, Theis FJ. SCANPY: large-scale single-cell gene expression data analysis. Genome Biol. 2018;19(1). doi:10.1186/s13059-017-1382-0

