## Supplementary Figures for "scMAGS: Marker gene selection from scRNA-seq data for spatial transcriptomics studies"

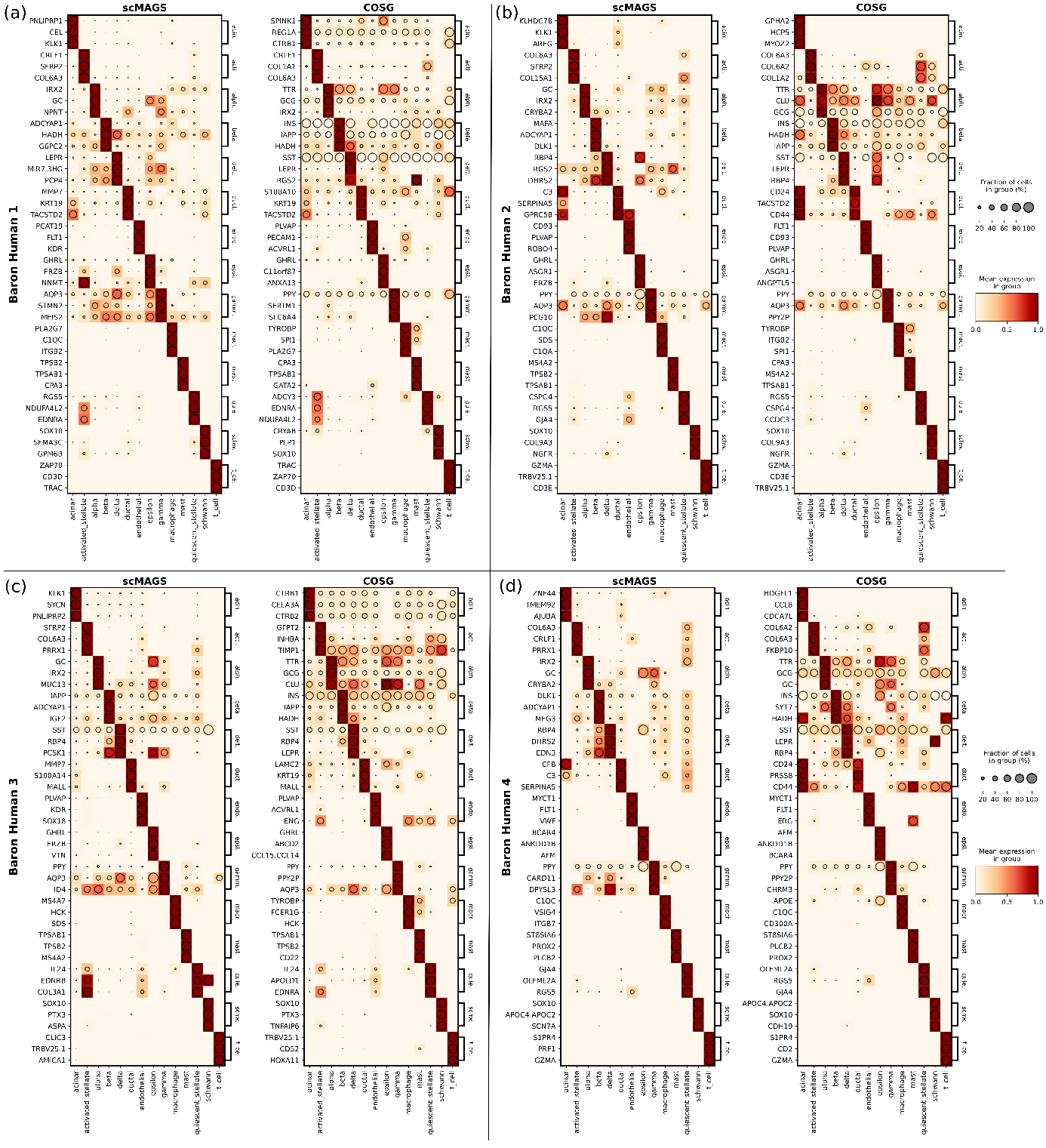

**Figure S1.** Dotplots obtained by the scMAGS and COSG methods for the **(a)** Baron Human 1 **(b)** Baron Human 2 **(c)** Baron Human 3 **(d)** Baron Human 4 datasets.

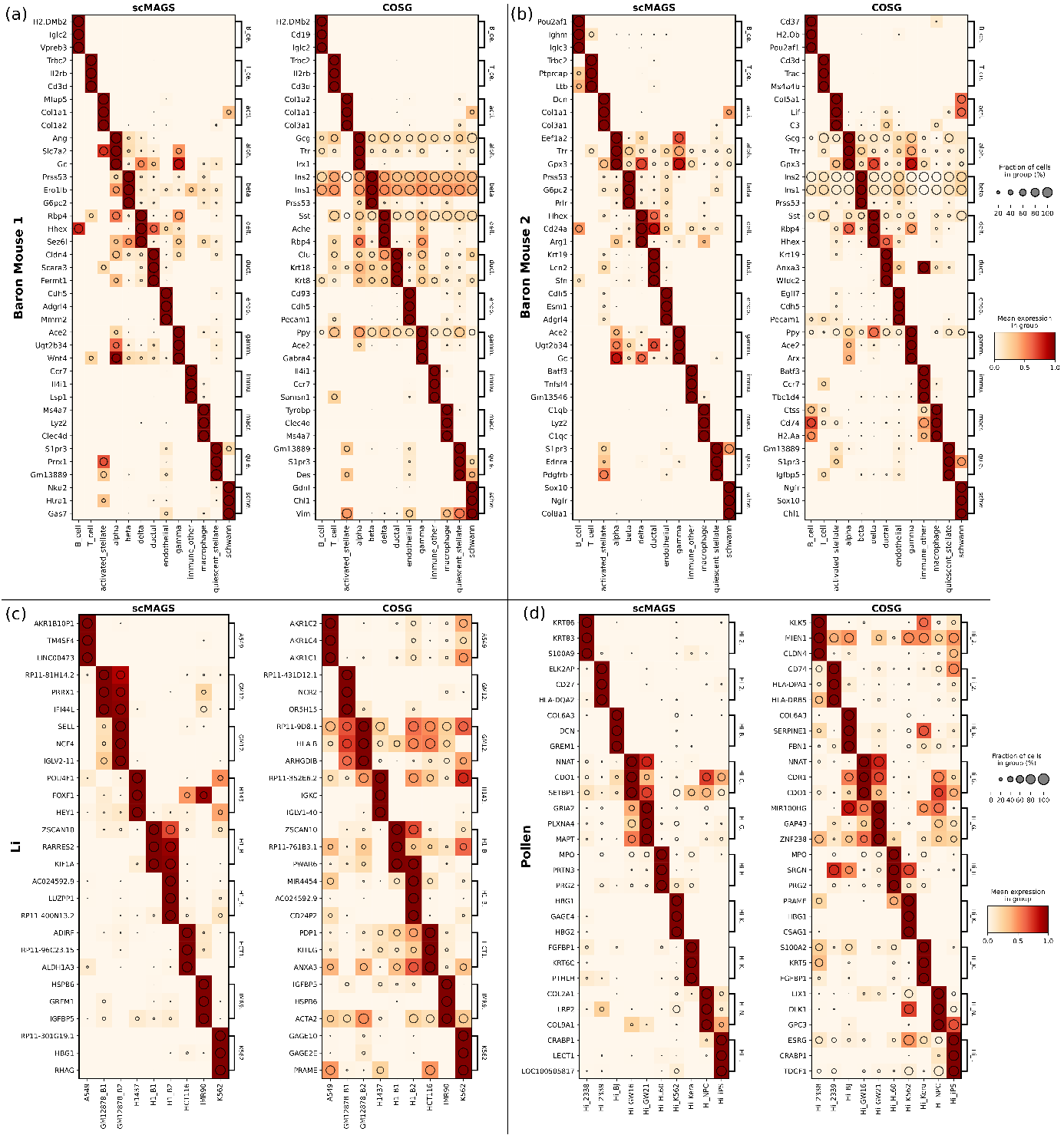

**Figure S2.** Dotplots obtained by the scMAGS and COSG methods for the **(a)** Baron Mouse 1 **(b)** Baron Mouse 2 **(c)** Li **(d)** Pollen datasets.

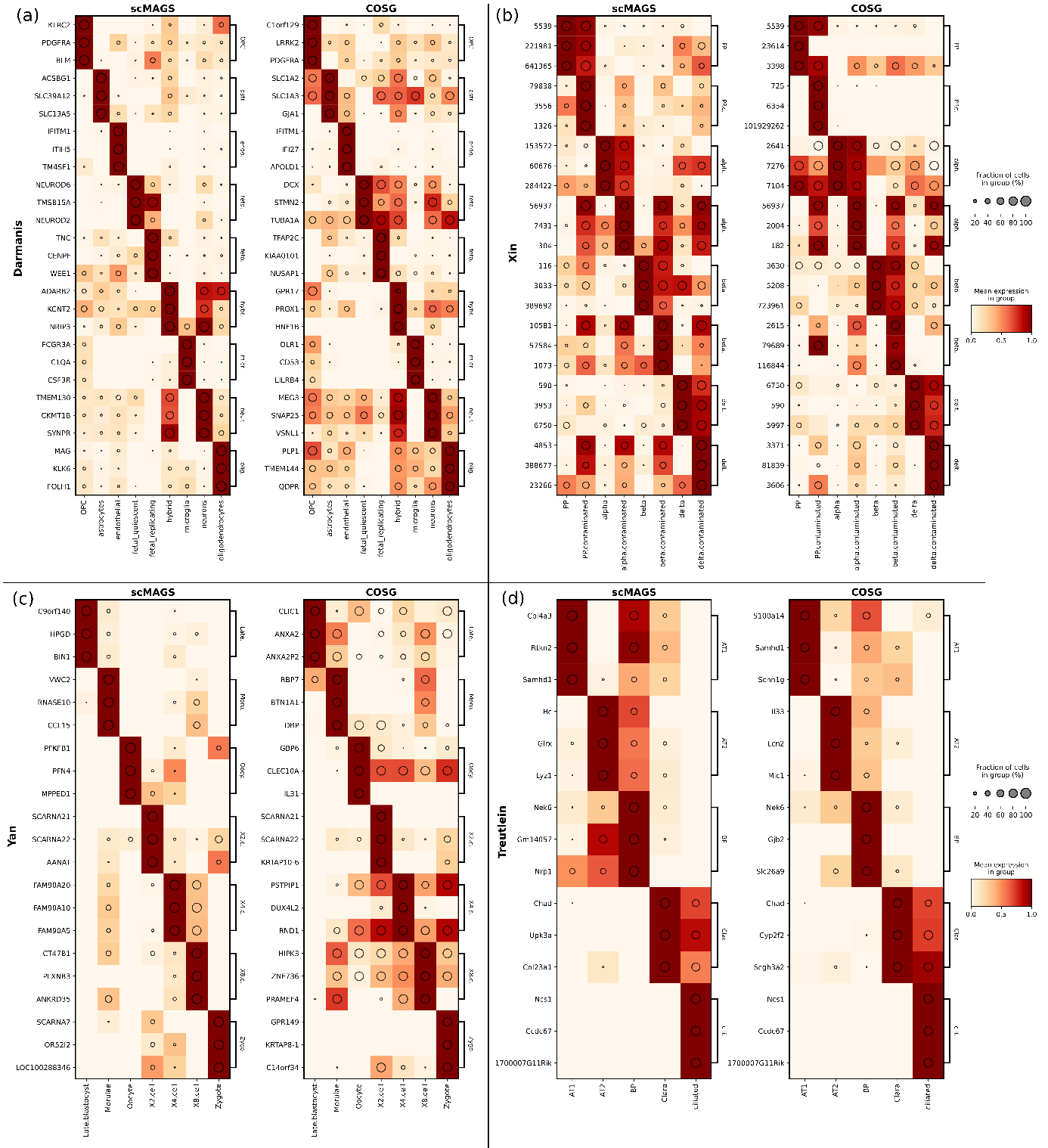

**Figure S3.** Dotplots obtained by the scMAGS and COSG methods for the **(a)** Darmanis **(b)** Xin **(c)** Yan **(d)** Treutlein datasets.

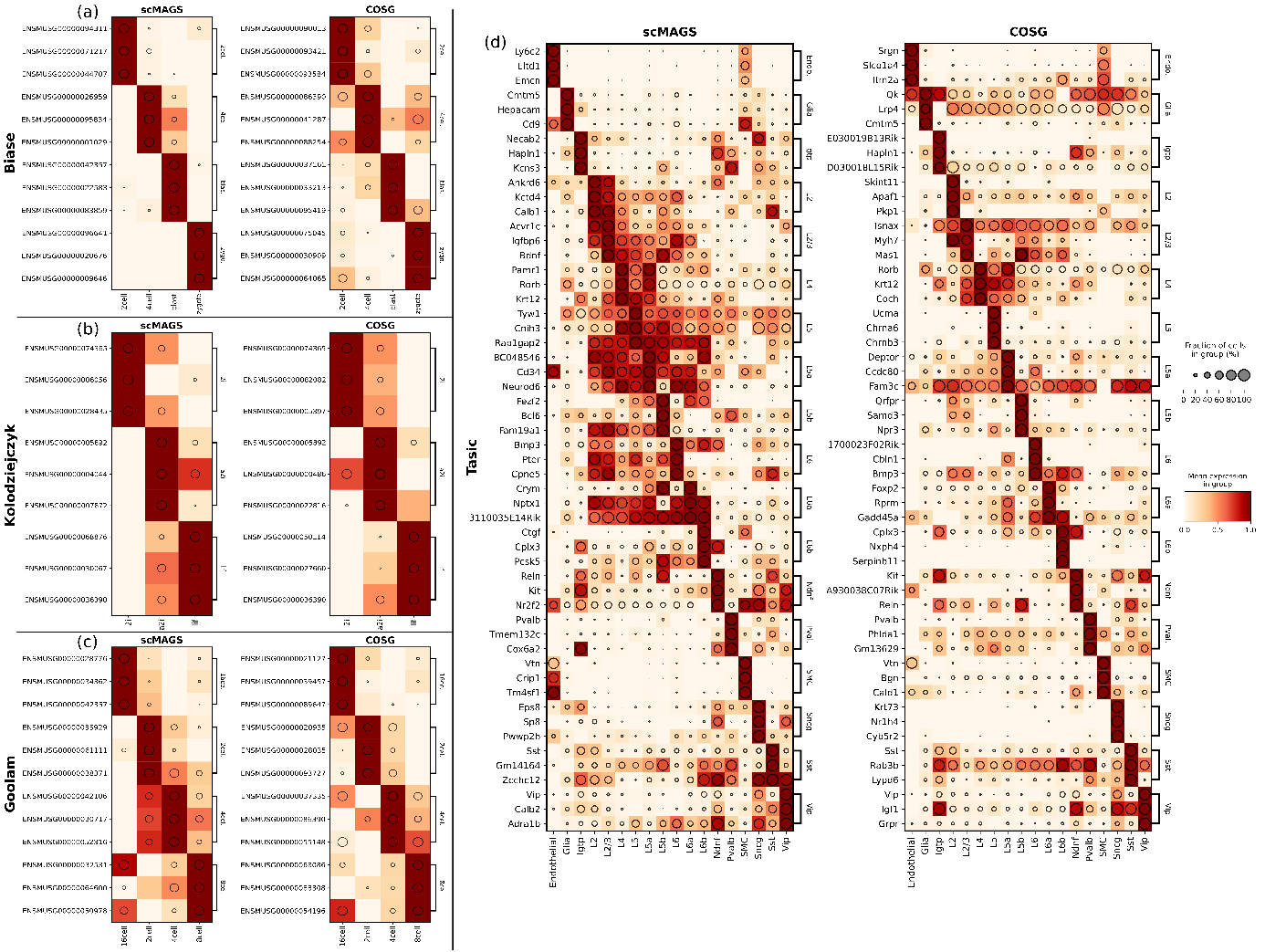

**Figure S4.** Dotplots obtained by the scMAGS and COSG methods for the **(a)** Biase **(b)** Kolodziejczyk **(c)** Goolam **(d)** Tasic datasets.

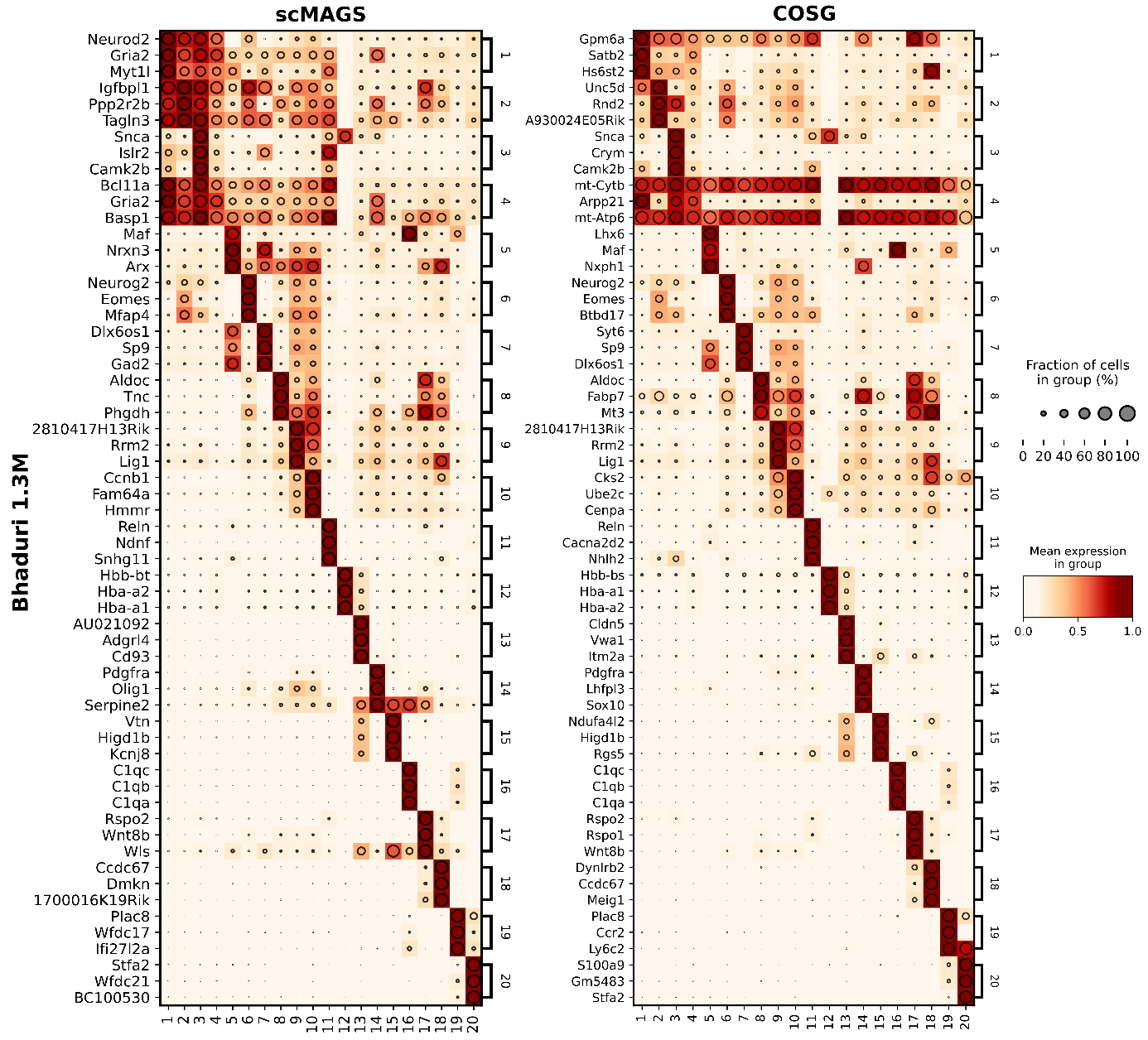

**Figure S5.** Dotplots obtained by the scMAGS and COSG methods for the Bhaduri dataset.
