## Supplementary Tables for "scMAGS: Marker gene selection from scRNA-seq data for spatial transcriptomics studies"

**Table S1.** Datasets used to evaluate the performance of the methods

| **Data-set** | **Number Of Cells** | **Number Of Genes** | **Number of Cell Types** | **Accession Code** |
| --- | --- | --- | --- | --- |
| Biase [1] | 56 | 25737 | 4 | [GSE57249](https://www.ncbi.nlm.nih.gov/geo/query/acc.cgi?acc=GSE57249) |
| Yan [2] | 90 | 20124 | 7 | [GSE36552](https://www.ncbi.nlm.nih.gov/geo/query/acc.cgi?acc=GSE36552) |
| Zeisel [3] | 3005 | 19972 | 9 | [GSE60361](https://www.ncbi.nlm.nih.gov/geo/query/acc.cgi?acc=GSE60361) |
| Li [4] | 561 | 55186 | 9 | [GSE81861](https://www.ncbi.nlm.nih.gov/geo/query/acc.cgi?acc=GSE81861) |
| Tasic [5] | 1679 | 24150 | 18 | [GSE71585](https://www.ncbi.nlm.nih.gov/geo/query/acc.cgi?acc=GSE71585) |
| Xin [6] | 1600 | 39851 | 8 | [GSE81608](https://www.ncbi.nlm.nih.gov/geo/query/acc.cgi?acc=GSE81608) |
| Darmanis [7] | 466 | 22088 | 9 | [GSE67835](https://www.ncbi.nlm.nih.gov/geo/query/acc.cgi?acc=GSE67835) |
| Baron Human 1 [8] | 1937 | 20125 | 14 | [GSE84133](https://www.ncbi.nlm.nih.gov/geo/query/acc.cgi?acc=GSE84133) |
| Baron Human 2 [8] | 1724 | 20125 | 14 | [GSE84133](https://www.ncbi.nlm.nih.gov/geo/query/acc.cgi?acc=GSE84133) |
| Baron Human 3 [8] | 3605 | 20125 | 14 | [GSE84133](https://www.ncbi.nlm.nih.gov/geo/query/acc.cgi?acc=GSE84133) |
| Baron Human 4 [8] | 1303 | 20125 | 14 | [GSE84133](https://www.ncbi.nlm.nih.gov/geo/query/acc.cgi?acc=GSE84133) |
| Baron Mouse 1 [8] | 822 | 14878 | 13 | [GSE84133](https://www.ncbi.nlm.nih.gov/geo/query/acc.cgi?acc=GSE84133) |
| Baron Mouse 2 [8] | 1064 | 14878 | 13 | [GSE84133](https://www.ncbi.nlm.nih.gov/geo/query/acc.cgi?acc=GSE84133) |
| Treutlein [9] | 80 | 23271 | 5 | [GSE52583](https://www.ncbi.nlm.nih.gov/geo/query/acc.cgi?acc=GSE52583) |
| Kolodziejczyk [10] | 704 | 38653 | 3 | [E-MTAB-2600](https://s3.amazonaws.com/scrnaseq-public-datasets/manual-data/kolodziejczyk/counttable_es.csv) |
| Goolam [11] | 124 | 41480 | 4 | [E-MTAB-3321](https://www.ebi.ac.uk/arrayexpress/files/E-MTAB-3321/E-MTAB-3321.processed.1.zip) |
| Pollen [12] | 301 | 23730 | 10 | [SRP041736](https://hemberg-lab.github.io/scRNA.seq.datasets/human/tissues/#pollen) |
| Kleshchevnikov [13] | 40532 | 31053 | 9 | [Sanger](https://cell2location.cog.sanger.ac.uk/browser.html) |
| Bhaduri [14] | 1.3 M | 27998 | 20 | [10X-Genomics](https://support.10xgenomics.com/single-cell-gene-expression/datasets/1.3.0/1M_neurons) |

**Table S2.** Computational time (in seconds) required by each method for each dataset

| **Data-Set** | **Number Of Cells** | **scmags** | **SMaSH** | **scGeneFit** | **COSG** |
| --- | --- | --- | --- | --- | --- |
| Biase | 56 | 4.035 | - | 1192.183 | 0.202 |
| Yan | 90 | 4.110 | - | 773.103 | 0.250 |
| Zeisel | 3005 | 6.132 | 76.998 | 745.508 | 0.762 |
| Li | 561 | 5.143 | - | - | 0.513 |
| Tasic | 1679 | 6.368 | - | - | 0.610 |
| Xin | 1600 | 5.784 | 65.968 | - | 0.815 |
| Darmanis | 466 | 4.360 | - | 9056.897 | 0.228 |
| Baron Human 1 | 1937 | 5.269 | - | 2411.984 | 0.608 |
| Baron Human 2 | 1724 | 4.992 | - | 1429.361 | 0.533 |
| Baron Human 3 | 3605 | 7.050 | - | 928.206 | 0.954 |
| Baron Human 4 | 1303 | 4.948 | - | 680.714 | 0.439 |
| Baron Mouse 1 | 822 | 3.869 | - | 575.568 | 0.286 |
| Baron Mouse 2 | 1064 | 4.309 | - | 401.690 | 0.309 |
| Treutlein | 80 | 3.986 | - | 2535.693 | 0.227 |
| Kolodziejczyk | 704 | 4.360 | 27.013 | 3332.770 | 0.385 |
| Goolam | 124 | 4.147 | - | 7325.235 | 0.368 |
| Pollen | 301 | 4.410 | - | - | 0.446 |
| Kleshchevnikov | 40532 | 159.897 | 912.117 | 4047.900 | 4.055 |
| Bhaduri | 1.3 M | 287.740 | - | - | 278.964 |

**Table S3.** Peak memory requirement (MB) of each method for each dataset

| **Data-Set** | **Number Of Cells** | **scmags** | **SMaSH** | **scGeneFit** | **COSG** |
| --- | --- | --- | --- | --- | --- |
| Biase | 56 | 201 | - | 35342 | 315 |
| Yan | 90 | 203 | - | 21940 | 326 |
| Zeisel | 3005 | 1267 | 6930 | 22299 | 2407 |
| Li | 561 | 704 | - | - | 1331 |
| Tasic | 1679 | 885 | - | - | 1704 |
| Xin | 1600 | 1431 | 7006 | - | 2508 |
| Darmanis | 466 | 353 | - | 26237 | 602 |
| Baron Human 1 | 1937 | 933 | - | 22328 | 1646 |
| Baron Human 2 | 1724 | 857 | - | 22366 | 1455 |
| Baron Human 3 | 3605 | 1495 | - | 22972 | 2812 |
| Baron Human 4 | 1303 | 672 | - | 22033 | 1129 |
| Baron Mouse 1 | 822 | 381 | - | 11974 | 666 |
| Baron Mouse 2 | 1064 | 523 | - | 12053 | 731 |
| Treutlein | 80 | 216 | - | 29099 | 329 |
| Kolodziejczyk | 704 | 739 | 3402 | 80312 | 1155 |
| Goolam | 124 | 271 | - | 92136 | 457 |
| Pollen | 301 | 294 | - | - | 557 |
| Kleshchevnikov | 40532 | 1854 | 122539 | 56460 | 3030 |
| Bhaduri | 1.3 M | 46949 | - | - | 112249 |

**References**

| 1. | Biase FH, Cao X, Zhong S. Cell fate inclination within 2-cell and 4-cell mouse embryos revealed by single-cell RNA sequencing. *Genome Res*. 2014;24(11):1787-1796. doi:10.1101/gr.177725.114 |
| --- | --- |
| 2. | Yan L, Yang M, Guo H, et al. Single-cell RNA-Seq profiling of human preimplantation embryos and embryonic stem cells. *Nat Struct Mol Biol*. 2013;20(9):1131-1139. doi:10.1038/nsmb.2660 |
| 3. | Zeisel A, Muñoz-Manchado AB, Codeluppi S, et al. Brain structure. Cell types in the mouse cortex and hippocampus revealed by single-cell RNA-seq. *Science*. 2015;347(6226):1138-1142. doi:10.1126/science.aaa1934 |
| 4. | Li H, Courtois ET, Sengupta D, et al. Reference component analysis of single-cell transcriptomes elucidates cellular heterogeneity in human colorectal tumors. *Nat Genet*. 2017;49(5):708-718. doi:10.1038/ng.3818 |
| 5. | Tasic B, Menon V, Nguyen TN, et al. Adult mouse cortical cell taxonomy revealed by single cell transcriptomics. *Nat Neurosci*. 2016;19(2):335-346. doi:10.1038/nn.4216 |
| 6. | Xin Y, Kim J, Okamoto H, et al. RNA sequencing of single human islet cells reveals type 2 diabetes genes. *Cell Metab*. 2016;24(4):608-615. doi:10.1016/j.cmet.2016.08.018 |
| 7. | Darmanis S, Sloan SA, Zhang Y, et al. A survey of human brain transcriptome diversity at the single cell level. *Proc Natl Acad Sci U S A*. 2015;112(23):7285-7290. doi:10.1073/pnas.1507125112 |
| 8. | Baron M, Veres A, Wolock SL, et al. A single-cell transcriptomic map of the human and mouse pancreas reveals inter- and intra-cell population structure. *Cell Syst*. 2016;3(4):346-360.e4. doi:10.1016/j.cels.2016.08.011 |
| 9. | Treutlein B, Brownfield DG, Wu AR, et al. Reconstructing lineage hierarchies of the distal lung epithelium using single-cell RNA-seq. *Nature*. 2014;509(7500):371-375. doi:10.1038/nature13173 |
| 10. | Kolodziejczyk AA, Kim JK, Tsang JCH, et al. Single cell RNA-sequencing of pluripotent states unlocks modular transcriptional variation. *Cell Stem Cell*. 2015;17(4):471-485. doi:10.1016/j.stem.2015.09.011 |
| 11. | Goolam M, Scialdone A, Graham SJL, et al. Heterogeneity in Oct4 and Sox2 targets biases cell fate in 4-cell mouse embryos. *Cell*. 2016;165(1):61-74. doi:10.1016/j.cell.2016.01.047 |
| 12. | Pollen AA, Nowakowski TJ, Shuga J, et al. Low-coverage single-cell mRNA sequencing reveals cellular heterogeneity and activated signaling pathways in developing cerebral cortex. *Nat Biotechnol*. 2014;32(10):1053-1058. doi:10.1038/nbt.2967 |
| 13. | Kleshchevnikov V, Shmatko A, Dann E, et al. Comprehensive mapping of tissue cell architecture via integrated single cell and spatial transcriptomics. *bioRxiv*. Published online 2020. doi:10.1101/2020.11.15.378125 |
| 14. | Bhaduri A, Nowakowski TJ, Pollen AA, Kriegstein AR. Identification of cell types in a mouse brain single-cell atlas using low sampling coverage. *BMC Biol*. 2018;16(1):113. doi:10.1186/s12915-018-0580-x |
